# Horizontal transfer and proliferation of Tsu4 in Saccharomyces paradoxus

**DOI:** 10.1101/287078

**Authors:** Casey M. Bergman

## Abstract

**Background:** Recent evidence suggests that horizontal transfer plays a significant role in the evolution of of transposable elements (TEs) in eukaryotes. Many cases of horizontal TE transfer (HTT) been reported in animals and plants, however surprisingly few examples of HTT have been reported in fungi.

**Findings:** Here I report evidence for a novel HTT event in fungi involving *Tsu4* in *Saccharomyces paradoxus* based on (i) high similarity between *Tsu4* elements in *S. paradoxus* and *S. uvarum*, (ii) a patchy distribution of *Tsu4* in *S. paradoxus* and general absence from its sister species *S. cerevisiae*, and (iii) discordance between the phylogenetic history of *Tsu4* sequences and species in the *Saccharomyces sensu stricto* group. Available data suggests the HTT event likely occurred somewhere in the Nearctic, Neotropic or Indo-Australian part of the *S. paradoxus* species range, and that a lineage related to *S. uvarum* or *S. eubayanus* was the donor species. The HTT event has led to massive proliferation of *Tsu4* in the South American lineage of *S. paradoxus*, which exhibits partial reproductive isolation with other strains of this species because of multiple reciprocal translocations. Full-length *Tsu4* elements are associated with both breakpoints of one of these reciprocal translocations.

**Conclusions:** This work shows that comprehensive analysis of TE sequences in essentially-complete genome assemblies derived from long-read sequencing provides new opportunities to detect HTT events in fungi and other organisms. This work also provides support for the hypothesis that HTT and subsequent TE proliferation can induce genome rearrangements that contribute to post-zygotic isolation in yeast.

## Main Text

Horizontal transfer is increasingly thought to play an important role in shaping the diversity of transposable elements (TEs) in eukaryotic genomes [1, 2, 3]. Since the initial discovery of horizontal transfer of the *P* element from *Drosophila willistoni* to *D. melanogaster* [4], a large number of cases of horizontal TE transfer (HTT) have been reported, especially among animals species (data compiled in [5]). However, surprisingly few cases of HTT have been reported in fungi [6, 7, 8, 9, 10, 11, 12], despite an abundance of genomic resources in this taxonomic group. Advances in long-read whole genome shotgun sequencing now allow comprehensive analysis of TE sequences in high-quality genome assemblies, and may therefore provide new opportunities for detecting HTT events in fungi and other organisms.

For example, a recent study by Yue *et al.* [13] reported essentially-complete PacBio genome assemblies for seven strains of *S. cerevisiae* and five strains of *S. paradoxus*. Analysis of TEs in these assemblies revealed a surprisingly high copy number for the *Ty4* family in one strain of *S. paradoxus* from South America (UFRJ50816; n=23 copies) [13]. This was a noteworthy observation for two reasons: (i) *Ty4* is typically found at low copy number in yeast strains [14, 15, 16], and (ii) S. American strains of *S. paradoxus* exhibit partial reproductive isolation with other strains of this species, which principally results from multiple reciprocal translocations thought to have arisen by unequal crossing-over between dispersed repetitive elements such as *Ty* elements [17].

I independently replicated the curious observation of exceptionally high *Ty4* copy number in *S. paradoxus* UFRJ50816 using a RepeatMasker-based annotation pipeline similar to that described in [11], which identifies and classifies Ty elements as full-length, truncated, or solo long terminal repeats (LTRs). Using the results of this initial annotation, I generated a multiple alignment of all full-length *Ty4* elements identified in these 12 assemblies. Preliminary phylogenetic analysis revealed that the full-length *Ty4* elements from *S. paradoxus* UFRJ50816 formed a monophyletic clade of very similar sequences that were highly divergent from other full-length *Ty4* elements identified in *S. cerevisiae* (S288c, Y12, YPS128) or *S. paradoxus* (N44). Surprisingly, BLAST analysis at NCBI using representative members of this divergent *Ty4*-like clade revealed that they were more similar to the *Tsu4* element from the related yeast species *S. uvarum* (Genbank: AJ439550) [18] than they were to the original *S. cerevisiae Ty4* query sequence (Genbank: S50671). This result suggested that the unusually high copy number of *Ty4* in *S. paradoxus* UFRJ50816 reported by Yue *et al.* [13] could actually be the consequence of rapid expansion of *Tsu4* following a HTT event from a *S. uvarum*-like donor.

To better characterize *Ty4* and *Tsu4* content in *S. cerevisiae* and *S. paradoxus*, I first identified a canonical *S. paradoxus Tsu4* element. To do this, I included the *S. uvarum Tsu4* query sequence in the TE library from [11] and re-annotated Ty elements in *S. paradoxus* UFRJ50816 using the same RepeatMasker-based strategy as above. I also performed *de novo* identification of full-length LTR elements in *S. paradoxus* UFRJ50816 using LTRharvest [19], then overlapped results from RepeatMasker and LTRharvest to identify full-length *Tsu4* elements, generated a consensus sequence from these elements, and finally identified the genomic copy (chrII:554570-560566) that clustered most closely with the consensus sequence of full-length *Tsu4* elements in a neighbor-joining tree. I then performed a final annotation of Ty elements in all 12 assemblies from Yue *et al.* [13] using the TE library from [11] plus the newly-identified *S. paradoxus Tsu4* canonical element. Full-length, truncated and solo LTR counts for *Tsu4, Ty4* and all other *Ty* families in these genomes can be found in Additional File 1; coordinates of annotated *Ty* elements in these genomes can be found in Additional File 2.

This improved annotation revealed solo LTRs for *Ty4* in all strains of *S. cerevisiae* and *S. paradoxus* with PacBio assemblies from from Yue *et al.* (Table 1). Solo LTRs arise by intra-element LTR-LTR recombination and serve as useful markers of past transpositional activity [20]. Additionally, *S. cerevisiae* S288c, Y12, and YPS128 as well as *S. paradoxus* N44 also contained a low copy number of full-length *Ty4* elements that is typical of this family. These results suggest that *Ty4* was present in the common ancestor of both *S. cerevisiae* and *S. paradoxus* and has been maintained at low copy number or become inactive in different lineages of each species.

**Table 1.**
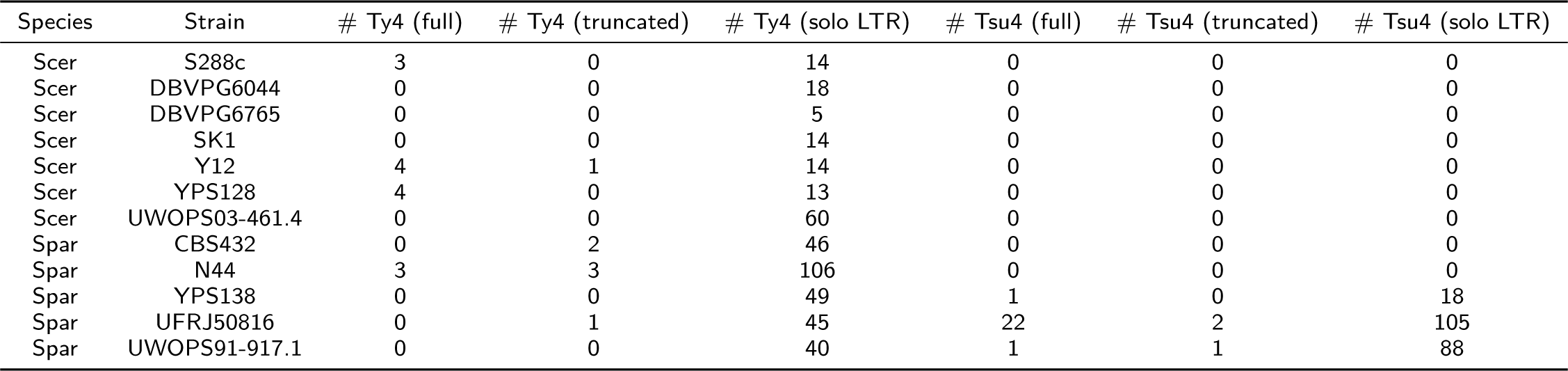
*Ty4* and *Tsu4* content in *S. cerevisiae* and *S. paradoxus* PacBio assemblies from Yue *et al.* [13]. Numbers of full-length elements (both LTRs present and internal region present with >95% coverage of canonical internal region), truncated elements (internal region present with *<*95% coverage of canonical internal region), or solo LTRs (no match to internal region) were estimated using a RepeatMasker (version 4.0.5) based strategy and a custom library of *Ty* elements from [11] supplemented with *S. paradoxus Tsu4*.

In contrast, I found evidence for full-length *Tsu4* sequences in only three strains of *S. paradoxus* from S. America (UFRJ50816, n=22), N. America (YPS138, n=1) and Hawaii (UWOPS91.917.1, n=1) (Table 1). The 22 full-length copies of *Tsu4* identified in UFRJ50816 are distributed on 11 different chromosomes and 12 of these full-length copies are flanked by 5-bp target site duplications that can be identified automatically by LTRharvest, suggesting proliferation in UFRJ50816 arose from active transposition and not simply duplication. *S. paradoxus* strains with full-length copies of *Tsu4* were devoid of full-length *Ty4* elements, and *vice versa*. Crucially, only these three *S. paradoxus* strains had solo LTRs for *Tsu4*, suggesting they were the only lineages in which *Tsu4* has been active in the past. The general absence of *Tsu4* in *S. cerevisiae* was confirmed by BLAST analysis of an additional 336 *S. cerevisiae* whole genome shotgun (WGS) assemblies at NCBI (taxid: 4932), which revealed only one nearly complete sequence with high similarity to *Tsu4* from a *S. cerevisiae* strain isolated from a rum distillery in the West Indies (>80% coverage and >80% identity, see below) [21]. The patchy distribution of active and relic *Tsu4* sequences in *S. paradoxus* and general absence from *S. cerevisiae* suggests that this element was not present in the common ancestor of these species, and was recently acquired by a *S. paradoxus* lineage somewhere in the Nearctic, Neotropic or Indo-Australian region, possibly in a strain lacking an active *Ty4*.

To provide further support for the hypothesis that *Tsu4* recently invaded *S. paradoxus* by HTT, I constructed a maximum likelihood phylogeny of all full-length *Ty4* and *Tsu4* sequences identified in the 12 strains of *S. cerevisiae* and *S. paradoxus* from Yue *et al.* [13] using RAxML [22]. In this analysis, I also included all complete or nearly-complete *Tsu4* elements identified by BLAST in 392 *Saccharomyces* WGS assemblies at NCBI (taxid: 4930) that had high similarity to the *S. uvarum Tsu4* query sequence (>80% coverage and >80% identity). These additional 12 *Tsu4* sequences include three sequences from the same strain of *S. uvarum* in which *Tsu4* was discovered, one sequence from *S. mikatae*, one sequence from *S. kudriavzevii*, one sequence each from four strains of *S. pastorianus*, two sequences from an unknown *Saccharomyces* species (strain M14) involved in lager brewing, and the single sequence from *S. cerevisiae* (strain 245) mentioned above (Additional File 3) [23, 21, 24, 25]. *S. mikatae* is the most closely related outgroup species to the *S. cerevisiae*/*S. paradoxus* clade, followed by *S. kudriavzevii*, then a clade containing *S. uvarum* and *S. eubayanus* (reviewed in [26]). *S. pastorianus* is a hybrid species used in lager brewing containing subgenomes from *S. cerevisiae* and *S. eubayanus* [27, 28, 24]. The multiple sequence alignment and maximum likelihood tree for this dataset can be found in Additional Files 4 and 5, respectively.

Figure 1A clearly shows that *Tsu4*-like sequences form a well-supported monophyletic clade that is distinct from the *Ty4* lineage present in *S. cerevisiae* and *S. paradoxus*. All *S. paradoxus Tsu4* sequences form a single clade that also contains the *Tsu4* sequence from *S. cerevisiae* strain 245, suggesting one initial HTT event into *S. paradoxus* followed by a secondary HTT event from *S. paradoxus* into *S. cerevisiae*. The most closely-related lineage to the *S. paradoxus Tsu4* clade is a clade containing sequences from *S. pastorianus* and *Saccharomyces sp. M14*, followed by a clade containing sequences from *S. uvarum*. The grouping of *Tsu4* sequences from the hybrid species *S. pastorianus* with those from *S. paradoxus* and *S. uvarum* can most parsimoniously be explained if *Tsu4* sequences from *S. pastorianus* are derived from the *S. eubayanus* component of the hybrid genome (see below). If this is true, then *Tsu4* in *S. paradoxus* could plausibly have arisen *via* HTT from *S. eubayanus*, the sister species to *S. uvarum. Tsu4* sequences from both *S. mikatae* and *S. kudriavzevii* are outgroups to the crown *Tsu4* lineage, but group more closely to the *Tsu4* lineage than to the *Ty4* lineage with strong support. The observation that *S. mikatae* groups more closely *Tsu4* with the crown *Tsu4* lineage than *S. kudriavzevii* is incompatible with the accepted species tree [26], suggesting unequal rates of evolution or another potential HTT event involving *Tsu4* between *S. mikatae* and the ancestor of *S. uvarum* and *S. eubayanus*. Despite unresolved issues with some aspects of the current *Tsu4* phylogeny, the fact that *S. uvarum* is the closest pure species clustering with the *S. paradoxus* clade is clearly incompatible with the accepted tree for these species [26] and this discordance provides support for the conclusion that *Tsu4* arose in *S. paradoxus* by HTT from *S. uvarum* or a closely related species like *S. eubayanus*.

**Figure 1.**
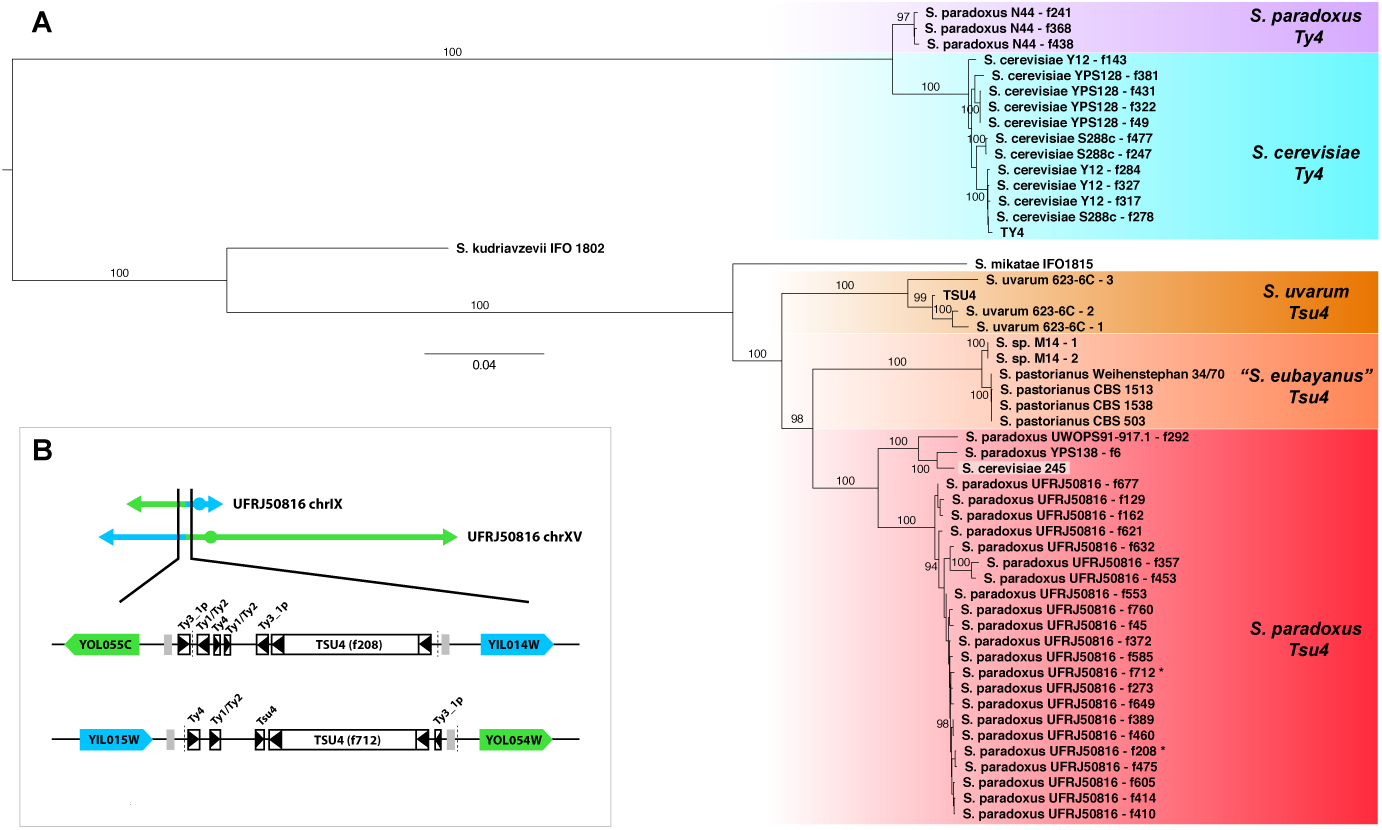
Evolution of the *Ty4* /*Tsu4* super-family in the *Saccharomyces sensu stricto* species group. **A**. Maximum likelihood phylogeny of all full-length *Ty4* and *Tsu4* elements from 12 strains of *S. cerevisiae* and *S. paradoxus* with PacBio assemblies from Yue *et al.* [13] plus all complete or nearly-complete *Tsu4* elements identified in *Saccharomyces* WGS assemblies at NCBI. Labels are shown for branches in the maximum likelihood tree that are supported by ≥90% of bootstrap replicates. The scale bar for branch lengths is in substitutions per site, and the tree is midpoint rooted. *Tsu4* sequences from hybrid strains (*S. pastorianus* and *Saccharomyces sp. M14*) were assigned to the *S. eubayanus* subgenome and presumed to have originated in *S. eubayanus*. **B**. Genome organization of the reciprocal translocation between chrIX and chrXV in UFRJ50816. Sequences from the standard arrangement chrIX are shown in blue, and sequences from the standard arrangement chrXV are shown in green. Protein-coding genes, tRNA genes, and solo LTRs are shown approximately to scale as solid arrows, grey rectangles and boxed arrowheads, respectively. Approximate translocation breakpoints in UFRJ50816 based on whole genome alignments can be localized to chrIX:252268-259232 and chrXV:320536-328356 (dashed lines). Full-length *Tsu4* elements are present in both translocation breakpoints. The *Tsu4* elements associated in the chrIX and chrXV reciprocal translocation between are denoted by asterices in panel **A**.

To address the origin of *Tsu4* sequences in *S. pastorianus* genomes and better understand potential donors for the HTT event, I aligned *S. pastorianus* WGS assemblies to a pan-genome comprised of *S. cerevisiae* S288c and *S. eubayanus* FM1318 [13, 28]. The *Tsu4* sequences from all four strains of *S. pastorianus* are contained on scaffolds that align best to scaffolds from the *S. eubayanus* subgenome (Additional File 6), consistent with the general lack of *Tsu4* in *S. cerevisiae* and the phylogenetic clustering of *S. pastorianus Tsu4* sequences with those from *S. uvarum*. In fact, *Tsu4* sequences for all four *S. pastorianus* strains align to the same location in *S. eubayanus* (scaffold NC 030972.1), which together with their tight clustering on the tree suggests they are alleles of the same insertion event. Likewise, alignment of the *Saccharomyces sp. M14* genome to a pan-genome of *S. eubayanus* and *S. cerevisiae* revealed large scaffolds that aligned to either scaffolds from *S. eubayanus* or chromosomes from *S. cerevisiae* (Additional File 7), in a similar manner to the *bona fide S. pastorianus* group 2/Frohberg strain W34/70 (Additional File 8). Thus, *Saccharomyces sp. M14* appears to be a hybrid of *S. cerevisiae* and *S. eubayanus* and may possibly be a previously-unidentified strain of *S. pastorianus*. The two *Tsu4* sequences from *Saccharomyces sp. M14* align to a different *S. eubayanus* scaffold (NC 030977.1) and form a cluster on the tree that is distinct from the *S. pastorianus Tsu4* sequences, suggesting they arose from different transposition events. None of the *S. pastorianus*/*Saccharomyces sp. M14 Tsu4* insertions are found in the *S. eubayanus* reference genome, nor are any other complete or nearly-complete *Tsu4* sequences. These results suggest that the *Tsu4* sequences in *S. pastorianus*/*Saccharomyces sp. M14* are derived from a *S. eubayanus* lineage with an active *Tsu4* element that is somewhat divergent from the *S. eubayanus* reference strain (from Northwestern Patagonia in Argentina [27]). Overall, the *S. eubayanus* subgenome localization and close affinity of *S. pastorianus*/*Saccharomyces sp. M14* with *S. paradoxus Tsu4* sequences suggests that a *S. eubayanus* lineage is a viable donor for the *Tsu4* element that invaded *S. paradoxus*.

Within *S. paradoxus*, the deepest well-supported branches in the *S. paradoxus Tsu4* clade are between N. American/Hawaiian and S. American *S. paradoxus* lineages, suggesting the HTT event predates separation of these lineages and could therefore have occurred anywhere in the ancestral range for *S. paradoxus* in the Nearctic, Neotropic or Indo-Australian regions. All of the *Tsu4* sequences in UFRJ50816 form a single clade, but bootstrap support for most branches within this clade are low, consistent with a recent proliferation event occurring after separation of S. American *S. paradoxus* from N. American and Hawaiian lineages. The single *S. cerevisiae Tsu4* sequence found in a strain 245 (isolated from the French West Indies) clusters strongly with the *Tsu4* sequence from the N. American *S. paradoxus* strain YPS138. Introgression of *S. paradoxus* DNA into *S. cerevisiae* has been observed previously [29, 30, 31, 32, 33], and thus introgression between a N. American-like lineage of *S. paradoxus Tsu4* and *S. cerevisiae* in the Caribbean could explain this secondary HTT event. As in N. American and Hawaiian lineages of *S. paradoxus, Tsu4* in this *S. cerevisiae* lineage has not led to widespread proliferation, suggesting the high copy number of *Tsu4* in UFRJ50816 is exceptional.

*S. paradoxus* UFRJ50816 was originally thought to represent a distinct species called *S. cariocanus* based on partial reproductive isolation with *S. paradoxus* tester strains [34, 35, 29]. At least five reciprocal translocations have been identified on the lineage leading to UFRJ50816 relative to the standard *S. paradoxus* karyotype that account for most of this reproductive isolation [17, 36, 13]. To test whether the recent proliferation of of *Tsu4* in UFRJ50816 has induced genome rearrangements involved in reproductive isolation, I identified translocation breakpoints in *S. paradoxus* UFRJ50816 relative to the standard karyotype *S. paradoxus* strain CBS432 using Mummer [37] and Ribbon [38]. Only one out of five translocations showed clear evidence for *Tsu4* sequences at both breakpoints. Intriguingly, both breakpoints of the translocation between chrIX and chrXV (which has recently been shown to reduce spore viability by approximately 50% [36]) each contained a full-length *Tsu4* element (Figure 1B). These two elements are from the major clade of *Tsu4* sequences found only in UFRJ50816, and are oriented in the directions expected if they were involved in a reciprocal exchange event. These results indicate that recent ectopic exchange among *Tsu4* sequences is not the primary cause of the majority of translocations in UFRJ50816, however *Tsu4* proliferation may have facilitated some genome rearrangements in the UFRJ50816 lineage.

In conclusion, here I report evidence for a novel HTT event in fungi involving *Tsu4* in *S. paradoxus* based on (i) high similarity between *Tsu4* elements in *S. paradoxus* and *S. uvarum*, (ii) a patchy distribution of *Tsu4* in *S. paradoxus* and general absence from its sister species *S. cerevisiae*, and (iii) discordance between the phylogenetic history of *Tsu4* sequences and host species trees. Using available genome assemblies, I conclude that the HTT event likely occurred in the Nearctic, Neotropic or Indo-Australian region, and that the donor species could be a lineage of either *S. uvarum* or *S. eubayanus*. This scenario is plausible since both *S. uvarum* and *S. eubayanus* have been sampled from sites in N. America and S. America that overlap or are in close proximity to the predicted range of *S. paradoxus* [27, 39, 40, 41, 42], and *S. uvarum* has been isolated from the same field sites as *S. paradoxus* in N. America [43]. Future work will hopefully refine the donor lineage and geographic location of the *Tsu4* HTT event, as well as the extent of the spread of *Tsu4* in *S. paradoxus*. I also show that full-length *Tsu4* elements are associated with the breakpoints of a reciprocal translocation that provides partial reproductive isolation between lineages of *S. paradoxus* from S. America and the rest of the world. These findings together with related work on *Ty2* in *S. cerevisiae* [8, 11, 44] provide support for the hypothesis that HTT and subsequent proliferation can induce genome rearrangements that contribute to post-zygotic isolation in yeast.

## List of abbreviations

TE: transposable element;
HTT: horizontal TE transfer;
LTR: long terminal repeat.

## Declarations

### Availability of data and materials

All data generated or analyzed during this study are included in this published article and its supplementary information files.

### Author’s contributions

CMB conceived of the project, performed the research, analyzed the data, and wrote the paper.

## Acknowledgements

CMB thanks Shan-ho Tsai and Yecheng Huang for bioinformatics application support; the Georgia Advanced Computing Resource Center for computing time; and Guilherme Dias and Douda Bensasson for comments on the manuscript.

## Additional Files

Additional File 1 — *Ty* content in *S. cerevisiae* and *S. paradoxus* PacBio assemblies

Numbers of full-length elements (both LTRs and internal region present with >95% coverage of canonical internal region), truncated elements (internal region present with *<*95% coverage of canonical internal region), or solo LTRs (no match to internal region) were identified using a RepeatMasker (version 4.0.5) based strategy and a custom library of *Ty* elements from [11] supplemented with *S. paradoxus Tsu4*.

Additional File 2 — Coordinates of *Ty* elements in *S. cerevisiae* and *S. paradoxus* PacBio assemblies.

Zip file of BED-formatted genome annotations for 12 strains of *S. cerevisiae* and *S. paradoxus* with PacBio assemblies. Full-length elements (f), truncated elements (t), or solo LTRs (s) were identified using a RepeatMasker (version 4.0.5) based strategy and a custom library of *Ty* elements from [11] supplemented with *S. paradoxus Tsu4*.

Additional File 3 — Summary of complete or nearly-complete *Tsu4* sequences identified in *Saccharomyces* whole genome assemblies at NCBI.

Accession number and coordinates of 12 complete or nearly-complete *Tsu4* elements identified by BLAST in 392 *Saccharomyces* (taxid: 4930) WGS assemblies at NCBI that had high similarity (>80% coverage and >80% identity) to the *Tsu4* query sequence (Genbank: AJ439550).

Additional File 4 — Multiple sequence alignment of *Ty4* and *Tsu4* elements in *Saccharomyces sensu stricto* species. Multiple sequence alignment of all full-length *Ty4* /*Tsu4* elements from 12 strains of *S. cerevisiae* and *S. paradoxus* with PacBio assemblies from Yue *et al.* [13] plus all complete or nearly-complete *Tsu4* elements identified in *Saccharomyces* WGS assemblies at NCBI (>80% coverage and >80% identity relative to the *Tsu4* query sequence from *S. uvarum*). Fasta files of *Ty4* /*Tsu4* sequences from all strains plus the *Ty4* and *Tsu4* query sequences were concatenated together and aligned using MAFFT (version 7.273-e; options: –thread 28) [45].

Additional File 5 — Maximum likelihood tree file for *Ty4* and *Tsu4* elements in *Saccharomyces sensu stricto* species. Newick-formatted file of the maximum-likelihood tree of all full-length *Ty4* /*Tsu4* elements from 16 strains of *S. cerevisiae* and *S. paradoxus* plus all complete or nearly-complete *Tsu4* elements identified in *Saccharomyces* WGS assemblies at NCBI. Maximum-likelihood phylogenetic analysis was performed on the multiple alignment in **Additional File 4** using RAxML (version: 8.2.4; options -T 28 -f a -x 12345 -p 12345 -N 100 -m GTRGAMMA) [22] excluding positions 1-166 and 6086-6476.

Additional File 6 — Coordinates of best-matches to scaffolds from *S. pastorianus* and *Saccharomyces sp. M14* containing *Tsu4* sequences *vs.* a pan-genome of *S. eubayanus* and *S. cerevisiae* genomes.

*S. pastorianus* and *Saccharomyces sp. M14* scaffolds were aligned to a pan-genome composed of scaffolds from *S. eubayanus* FM1318 (Genbank: GCF 001298625.1) and chromosomes from *S. cerevisiae* S288c (from [13]). Alignments were generated using nucmer (default parameters), delta-filter (options: -1 -l 2000), and show-coords (options: -lTH) in mummer 3.23 [37].

Additional File 7 — Dot-plot of the Chinese lager strain *Saccharomyces sp. M14 vs.* a pan-genome of *S. eubayanus* and *S. cerevisiae* genomes.

Dot-plot of *Saccharomyces sp. M14* scaffolds aligned to a pan-genome composed of scaffolds from *S. eubayanus* FM1318 (Genbank: GCF 001298625.1) and chromosomes from *S. cerevisiae* S288c (from [13]), showing that *Saccharomyces sp. M14* contains subgenomes from both species and that this strain may be a previously-unidentified strain of the lager brewing species *S. pastorianus*. The dot-plot was generated using nucmer (default parameters) and mummerplot (options: --size large -fat --color -f --png) in mummer 3.23 [37].

Additional File 8 — Dot-plot of the *S. pastorianus* group 2/Frohberg strain W34/70 *vs.* a pan-genome of *S. eubayanus* and *S. cerevisiae* genomes.

Dot-plot of *S. pastorianus* group 2/Frohberg strain W34/70 scaffolds aligned to a pan-genome composed of scaffolds from *S. eubayanus* (Genbank: GCF 001298625.1) and chromosomes from *S. cerevisiae* (S288c from [13]), showing that *S. pastorianus* group 2/Frohberg strain W34/70 contains subgenomes from both species in a similar pattern as for *Saccharomyces sp. M14* (see **Additional File 7**). The dot-plot was generated using nucmer (default parameters) and mummerplot (options: --size large -fat --color -f --png) in mummer 3.23 [37]

## References

1. Schaack, S., Gilbert, C., Feschotte, C.: Promiscuous DNA: horizontal transfer of transposable elements and why it matters for eukaryotic evolution. Trends Ecol Evol (Amst) 25(9), 537–546 (2010). doi:10.1016/j.tree.2010.06.001

2. Wallau, G.L., Ortiz, M.F., Loreto, E.L.S.: Horizontal transposon transfer in eukarya: detection, bias, and perspectives. Genome Biol Evol 4(8), 801–811 (2012). doi:10.1093/gbe/evs055. Accessed 2018-02-07

3. Wallau, G.L., Vieira, C., Loreto, E.L.S.: Genetic exchange in eukaryotes through horizontal transfer: connected by the mobilome. Mobile DNA 9, 6 (2018). doi:10.1186/s13100-018-0112-9. Accessed 2018-02-07

4. Daniels, S.B., Peterson, K.R., Strausbaugh, L.D., Kidwell, M.G., Chovnick, A.: Evidence for horizontal transmission of the P transposable element between Drosophila species. Genetics 124(2), 339–55 (1990)

5. Dotto, B.R., Carvalho, E.L., Silva, A.F., Silva, D., Fernando, L., Pinto, P.M., Ortiz, M.F., Wallau, G.L.: WHTT-DB: Horizontally transferred transposable elements database. Bioinformatics 31(17), 2915–2917 (2015). doi:10.1093/bioinformatics/btv281. Accessed 2018-02-07

6. Dobinson, K.F., Harris, R.E., Hamer, J.E.: Grasshopper, a long terminal repeat (LTR) retroelement in the phytopathogenic fungus Magnaporthe grisea. Mol Plant Microbe Interact 6(1), 114–126 (1993)

7. Daboussi, M.-J., Daviére, J.-M., Graziani, S., Langin, T.: Evolution of the Fot1 transposons in the genus Fusarium: discontinuous distribution and epigenetic inactivation. Mol Biol Evol 19(4), 510–520 (2002). doi:101093/oxfordjournalsmolbeva004106

8. Liti, G., Peruffo, A., James, S.A., Roberts, I.N., Louis, E.J.: Inferences of evolutionary relationships from a population survey of LTR-retrotransposons and telomeric-associated sequences in the Saccharomyces sensu stricto complex. Yeast 22(3), 177–92 (2005). Liti, Gianni Peruffo, Antonella James, Steve A Roberts, Ian N Louis, Edward J Research Support, Non-U.S. Gov’t England Yeast (Chichester, England) Yeast. 2005 Feb;22(3):177–92.

9. Novikova, O., Fet, V., Blinov, A.: Non-LTR retrotransposons in fungi. Funct Integr Genomics 9(1), 27–42 (2009). doi:10.1007/s10142-008-0093-8. Accessed 2018-02-07

10. Amyotte, S.G., Tan, X., Pennerman, K., del Mar Jimenez-Gasco, M., Klosterman, S.J., Ma, L.-J., Dobinson, K.F., Veronese, P.: Transposable elements in phytopathogenic Verticillium spp.: insights into genome evolution and inter- and intra-specific diversification. BMC Genomics 13, 314 (2012). doi:10.1186/1471-2164-13-314. Accessed 2018-02-07

11. Carr, M., Bensasson, D., Bergman, C.M.: Evolutionary genomics of transposable elements in Saccharomyces cerevisiae. PloS One 7(11), 50978 (2012). doi:101371/journalpone0050978. Accessed 2012-12-03

12. Sarilar, V., Bleykasten-Grosshans, C., Neuvéglise, C.: Evolutionary dynamics of hAT DNA transposon families in Saccharomycetaceae. Genome Biol Evol 7(1), 172–190 (2015). doi:10.1093/gbe/evu273. Accessed 2018-02-07

13. Yue, J.-X., Li, J., Aigrain, L., Hallin, J., Persson, K., Oliver, K., Bergström, A., Coupland, P., Warringer, J., Lagomarsino, M.C., Fischer, G., Durbin, R., Liti, G.: Contrasting evolutionary genome dynamics between domesticated and wild yeasts. Nat Genet 49(6), 913–924 (2017). doi:10.1038/ng.3847. Accessed 2017-07-27

14. Stucka, R., Lochmäller, H., Feldmann, H.: Ty4, a novel low-copy number element in Saccharomyces cerevisiae: one copy is located in a cluster of Ty elements and tRNA genes. Nucleic Acids Res 17(13), 4993–5002 (1989). doi:10.1093/nar/17.13.4993. Accessed 2015-09-15

15. Kim, J.M., Vanguri, S., Boeke, J.D., Gabriel, A., Voytas, D.F.: Transposable elements and genome organization: a comprehensive survey of retrotransposons revealed by the complete Saccharomyces cerevisiae genome sequence. Genome Research 8(5), 464–78 (1998)

16. Bleykasten-Grosshans, C., Friedrich, A., Schacherer, J.: Genome-wide analysis of intraspecific transposon diversity in yeast. BMC Genomics 14, 399 (2013). doi:10.1186/1471-2164-14-399

17. Fischer, G., James, S.A., Roberts, I.N., Oliver, S.G., Louis, E.J.: Chromosomal evolution in Saccharomyces. Nature 405(6785), 451–4 (2000). doi:10.1038/35013058

18. Neuvéglise, C., Feldmann, H., Bon, E., Gaillardin, C., Casaregola, S.: Genomic evolution of the long terminal repeat retrotransposons in hemiascomycetous yeasts. Genome Research 12(6), 930–43 (2002). doi:10.1101/gr.219202

19. Ellinghaus, D., Kurtz, S., Willhoeft, U.: LTRharvest, an efficient and flexible software for de novo detection of LTR retrotransposons. BMC Bioinformatics 9(1), 18 (2008). doi:10.1186/1471-2105-9-18

20. Fink, G., Boeke, J., Garfinkel, D.: The mechanism and consequences of retrotransposition. Trends in Genetics 2, 118–123 (1986). doi:DOI: 10.1016/0168-9525(86)90200-3

21. Marsit, S., Mena, A., Bigey, F., Sauvage, F.-X., Couloux, A., Guy, J., Legras, J.-L., Barrio, E., Dequin, S., Galeote, V.: Evolutionary advantage conferred by an eukaryote-to-eukaryote gene transfer event in wine yeasts. Mol Biol Evol 32(7), 1695–1707 (2015). doi:10.1093/molbev/msv057. Accessed 2018-02-20

22. Stamatakis, A.: RAxML version 8: a tool for phylogenetic analysis and post-analysis of large phylogenies. Bioinformatics 30(9), 1312–1313 (2014). doi:10.1093/bioinformatics/btu033

23. Cliften, P., Sudarsanam, P., Desikan, A., Fulton, L., Fulton, B., Majors, J., Waterston, R., Cohen, B.A., Johnston, M.: Finding functional features in Saccharomyces genomes by phylogenetic footprinting. Science 301(5629), 71–76 (2003). doi:10.1126/science.1084337

24. Okuno, M., Kajitani, R., Ryusui, R., Morimoto, H., Kodama, Y., Itoh, T.: Next-generation sequencing analysis of lager brewing yeast strains reveals the evolutionary history of interspecies hybridization. DNA Res 23(1), 67–80 (2016). doi:10.1093/dnares/dsv037. Accessed 2018-02-20

25. Liu, C., Li, Q., Niu, C., Zheng, F., Li, Y., Zhao, Y., Yin, X.: Genome sequence of the lager-brewing yeast Saccharomyces sp. strain M14, used in the high-gravity brewing industry in China. Genome Announc 5(43) (2017). doi:10.1128/genomeA.01194-17. Accessed 2018-02-20

26. Borneman, A.R., Pretorius, I.S.: Genomic insights into the Saccharomyces sensu stricto complex. Genetics 199(2), 281–291 (2015). doi:10.1534/genetics.114.173633. Accessed 2018-02-19

27. Libkind, D., Hittinger, C.T., Valerio, E., Goncalves, C., Dover, J., Johnston, M., Goncalves, P., Sampaio, J.P.: Microbe domestication and the identification of the wild genetic stock of lager-brewing yeast. Proc Natl Acad Sci USA 108(35), 14539–14544 (2011). doi:10.1073/pnas.1105430108. Accessed 2018-02-13

28. Baker, E., Wang, B., Bellora, N., Peris, D., Hulfachor, A.B., Koshalek, J.A., Adams, M., Libkind, D., Hittinger, C.T.: The genome sequence of Saccharomyces eubayanus and the domestication of lager-brewing yeasts. Mol Biol Evol 32(11), 2818–2831 (2015). doi:10.1093/molbev/msv168. Accessed 2017-08-22

29. Liti, G., Barton, D.B.H., Louis, E.J.: Sequence diversity, reproductive isolation and species concepts in Saccharomyces. Genetics 174(2), 839–850 (2006). doi:10.1534/genetics.106.062166

30. Doniger, S.W., Kim, H.S., Swain, D., Corcuera, D., Williams, M., Yang, S.-P., Fay, J.C.: A catalog of neutral and deleterious polymorphism in yeast. PLoS Genet 4(8), 1000183 (2008). doi:101371/journalpgen1000183

31. Muller, L.A.H., McCusker, J.H.: A multi-species based taxonomic microarray reveals interspecies hybridization and introgression in Saccharomyces cerevisiae. FEMS Yeast Res 9(1), 143–152 (2009). doi:10.1111/j.1567-1364.2008.00464.x. Accessed 2018-02-13

32. Strope, P.K., Skelly, D.A., Kozmin, S.G., Mahadevan, G., Stone, E.A., Magwene, P.M., Dietrich, F.S., McCusker, J.H.: The 100-genomes strains, an S. cerevisiae resource that illuminates its natural phenotypic and genotypic variation and emergence as an opportunistic pathogen. Genome Res 25(5), 762–774 (2015). doi:10.1101/gr.185538.114. Accessed 2015-10-22

33. Almeida, P., Barbosa, R., Bensasson, D., Goncalves, P., Sampaio, J.P.: Adaptive divergence in wine yeasts and their wild relatives suggests a prominent role for introgressions and rapid evolution at noncoding sites. Mol Ecol 26(7), 2167–2182 (2017). doi:10.1111/mec.14071

34. Naumov, G.I., Naumova, E.S., Hagler, A.N., Mendonça-Hagler, L.C., Louis, E.J.: A new genetically isolated population of the Saccharomyces sensu stricto complex from Brazil. Antonie van Leeuwenhoek 67(4), 351–355 (1995). doi:10.1007/BF00872934. Accessed 2018-02-21

35. Naumov, G.I., James, S.A., Naumova, E.S., Louis, E.J., Roberts, I.N.: Three new species in the Saccharomyces sensu stricto complex: Saccharomyces cariocanus, Saccharomyces kudriavzevii and Saccharomyces mikatae. Int J Syst Evol Microbiol 50 Pt 5, 1931–1942 (2000). doi:10.1099/00207713-50-5-1931

36. Almutawa, Q.: Impact of Chromosomal Translocations (CTs) on reproductive isolation and fitness in natural yeast isolates. PhD thesis, University of Manchester (2016). https://www.escholar.manchester.ac.uk/item/?pid=uk-ac-man-scw:307192 Accessed 2018-02-22

37. Kurtz, S., Phillippy, A., Delcher, A.L., Smoot, M., Shumway, M., Antonescu, C., Salzberg, S.L.: Versatile and open software for comparing large genomes. Genome Biol 5(2), 12 (2004). doi:10.1186/gb-2004-5-2-r12

38. Nattestad, M., Chin, C.-S., Schatz, M.C.: Ribbon: Visualizing complex genome alignments and structural variation. bioRxiv, 082123 (2016). doi:10.1101/082123. Accessed 2018-02-23

39. Almeida, P., Goncalves, C., Teixeira, S., Libkind, D., Bontrager, M., Masneuf-Pomarede, I., Albertin, W., Durrens, P., Sherman, D.J., Marullo, P., Todd Hittinger, C., Goncalves, P., Sampaio, J.P.: A Gondwanan imprint on global diversity and domestication of wine and cider yeast Saccharomyces uvarum. Nature Communications 5, 4044 (2014). doi:10.1038/ncomms5044. Accessed 2018-02-13

40. Peris, D., Sylvester, K., Libkind, D., Gonçalves, P., Sampaio, J.P., Alexander, W.G., Hittinger, C.T.: Population structure and reticulate evolution of Saccharomyces eubayanus and its lager-brewing hybrids. Mol Ecol 23(8), 2031–2045 (2014). doi:10.1111/mec.12702

41. Peris, D., Langdon, Q.K., Moriarty, R.V., Sylvester, K., Bontrager, M., Charron, G., Leducq, J.-B., Landry, C.R., Libkind, D., Hittinger, C.T.: Complex ancestries of lager-brewing hybrids were shaped by standing variation in the wild yeast Saccharomyces eubayanus. PLoS Genet 12(7), 1006155 (2016). doi:101371/journalpgen1006155. Accessed 2016-07-09

42. Robinson, H.A., Pinharanda, A., Bensasson, D.: Summer temperature can predict the distribution of wild yeast populations. Ecol Evol 6(4), 1236–1250 (2016). doi:10.1002/ece3.1919. Accessed 2018-03-05

43. Sampaio, J.P., Goncalves, P.: Natural populations of Saccharomyces kudriavzevii in Portugal are associated with oak bark and are sympatric with S. cerevisiae and S. paradoxus. Appl Environ Microbiol 74(7), 2144–2152 (2008). doi:10.1128/AEM.02396-07. Accessed 2018-02-19

44. Hou, J., Friedrich, A., de Montigny, J., Schacherer, J.: Chromosomal rearrangements as a major mechanism in the onset of reproductive isolation in Saccharomyces cerevisiae. Curr Biol 24(10), 1153–1159 (2014). doi:10.1016/j.cub.2014.03.063

45. Katoh, K., Kuma, K., Toh, H., Miyata, T.: MAFFT version 5: improvement in accuracy of multiple sequence alignment. Nucleic Acids Res 33(2), 511–8 (2005). 1362-4962 (Electronic) Evaluation Studies Journal Article

