## Supplementary figures and images for "Horizontal transfer and proliferation of Tsu4 in Saccharomyces paradoxus"

### Supplementary Materials

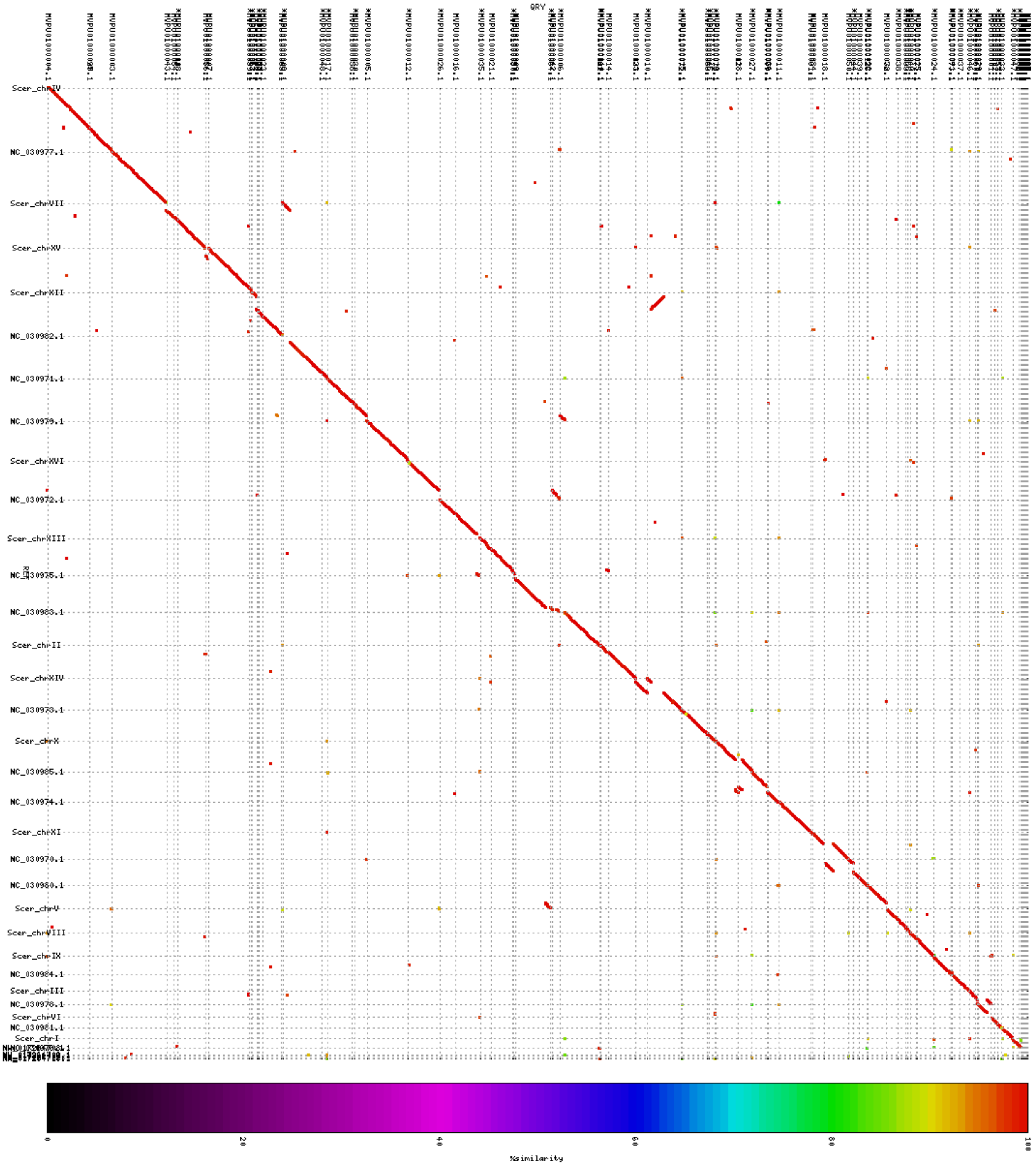

### Supplementary Materials

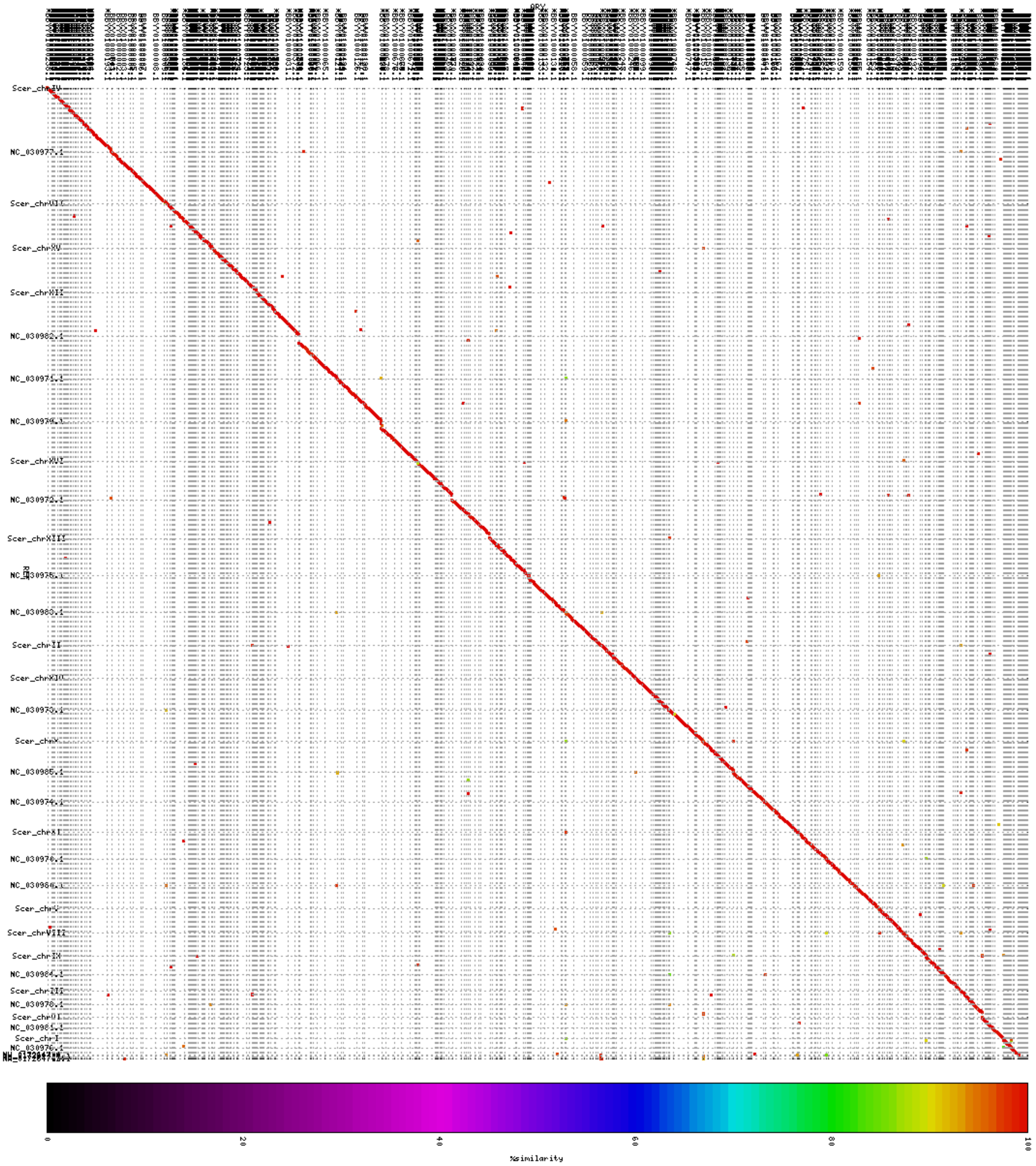
